# Methylthioadenosine reduces host inflammatory response by suppressing *Salmonella* virulence

**DOI:** 10.1101/287052

**Authors:** Jeffrey S. Bourgeois, Daoguo Zhou, Teresa L. M. Thurston, James J. Gilchrist, Dennis C. Koa

**Author notes:** JJG and DCK contributed equally to this work.

## Abstract

In order to deploy virulence factors at appropriate times and locations, microbes must rapidly sense and respond to various metabolite signals. Previously we showed transient elevation of the methionine-derived metabolite methylthioadenosine (MTA) in serum during systemic *Salmonella enterica* serovar Typhimurium (*S.* Typhimurium) infection. Here we explored the functional consequences of increased MTA concentrations on *S.* Typhimurium virulence. We found that MTA—but not other related metabolites involved in polyamine synthesis and methionine salvage—reduced motility, host cell pyroptosis, and cellular invasion. Further, we developed a genetic model of increased bacterial endogenous MTA production by knocking out the master repressor of the methionine regulon, *metJ*. Like MTA treated *S.* Typhimurium, the Δ*metJ* mutant displayed reduced motility, host cell pyroptosis, and invasion. These phenotypic effects of MTA correlated with suppression of flagellar and *Salmonella* pathogenicity island-1 (SPI-1) networks. Δ*metJ S.* Typhimurium had reduced virulence in oral infection of C57BL/6 mice. Finally, Δ*metJ* bacteria induced a less severe inflammatory cytokine response in a mouse sepsis model. These data provide a possible bacterial mechanism for our previous findings that pretreating mice with MTA dampens inflammation and prolongs survival. Together, these data indicate that exposure of *S.* Typhimurium to MTA or disruption of the bacterial methionine metabolism pathway is sufficient to suppress SPI-1 mediated processes, motility, and *in vivo* virulence.

**Significance:** *Salmonella enterica* serovar Typhimurium (*S*. Typhimurium) is a leading cause of gastroenteritis and bacteremia worldwide. Widespread multi-drug resistance, inadequate diagnostics, and the absence of a vaccine for use in humans, all contribute to the global burden of morbidity and mortality associated with *S.* Typhimurium infection. Here we find that increasing the concentration of the methionine derived metabolite methylthioadenosine, either in *S.* Typhimurium or in its environment, is sufficient to suppress virulence processes. These findings could be leveraged to inform future therapeutic interventions against *S.* Typhimurium aimed at manipulating either host or pathogen methylthioadenosine production.

## Introduction

Microbial communities within a mammalian host are bombarded by an array of intercellular, interspecies, and cross-kingdom metabolites and proteins. Cross-kingdom signaling plays important roles in *Salmonella* pathogenesis. For instance, in order to invade non-phagocytic host cells, *Salmonella* must deploy a secretion system encoded the *Salmonella* Pathogenicity Island-1 (SPI-1) (1, 2), which is regulated by many signals such as pH, bile, and short chain fatty acids (3–7). Together, these factors spatially limit the bacteria so that most invasion occurs in the ileum (8). Furthermore, recent work demonstrates that a host mimic of the bacterial AI-2 quorum molecule can directly impact *S*. Typhimurium gene expression *in vitro* by activating the *lsr* operon (9). Understanding how the bacteria’s environment influences *Salmonella* pathogenesis is important as it could help to inform future therapeutic interventions to suppress virulence.

One signal that may facilitate cross-talk between host and pathogen during infection is methylthioadenosine (MTA), a key metabolite in methionine metabolism. In addition to its role in protein synthesis, methionine is used in both eukaryotic and prokaryotic systems to generate S-Adenosyl methionine (SAM), which is a critical methyl donor for a number of reactions (10, 11). SAM catabolism results in a number of metabolic byproducts, including MTA and S-adenosylhomocysteine (SAH). In many eukaryotic and prokaryotic systems, MTA is recycled back into methionine, however, *E. coli* and *S*. Typhimurium cannot salvage methionine from MTA (12). Instead, *E. coli* and *Salmonella* spp. regulate intracellular MTA concentrations by using an MTA/SAH nucleosidase (*pfs)* to cleave MTA into 5’methythioribose and excreting it (13, 14). MTA regulation is considered to be critical for the bacterial cell as deletion of *pfs* impairs growth (15), but the effects of MTA on *Salmonella* virulence remain unknown.

Previously, our lab determined that MTA plays a multifaceted role in *Salmonella* infection. We originally identified MTA as a positive regulator of host cell pyroptosis, a rapid, proinflammatory form of cell death, during *Salmonella* infection (16). More recently, we showed that host MTA is released into plasma during *S*. Typhimurium infection, and that high plasma MTA levels are associated with poor sepsis outcomes in humans (17). Paradoxically, we showed that exogenous treatment with MTA suppressed sepsis-associated cytokines and extended the lifespan of mice infected with a lethal dose of *S*. Typhimurium (17). While consistent with previous reports that MTA acts as an anti-inflammatory molecule (18–20), this was in contrast to our findings that MTA primes cells to undergo pyroptosis. Together, these data led us to hypothesize that increased extracellular concentrations of MTA could potentially have independent effects on both the host and pathogen during infection.

Here we show that fluctuations in MTA levels regulate *S.* Typhimurium virulence *in vitro* and *in vivo*. Treatment of *S.* Typhimurium with exogenous MTA prior to infection or increasing endogenous bacterial production of MTA through genetic deletion of the methionine regulon suppressor, *metJ,* reduced the induction of pyroptosis and invasion *in vitro*. Furthermore, we report that both Δ*metJ* mutants and MTA treated bacteria demonstrate transcriptional, translational, and functional reductions in SPI-1 activity and motility. Finally, we find that Δ*metJ* mutants have reduced virulence *in vivo* and that disrupting the methionine metabolism pathway in the bacteria can influence the inflammatory state of the host. Together, these data reveal the importance of MTA and bacterial methionine metabolism for regulating *S.* Typhimurium virulence and host inflammation, and provide a possible example of host-pathogen metabolite cross-talk during infection.

## Results

### Exogenous MTA reduces the ability for *S.* Typhimurium to induce pyroptosis and invade host cells *in vitro*.

Previously, we demonstrated elevated concentrations of MTA in plasma during systemic infection of mice with *S.* Typhimurium (17). While effects of MTA on the host inflammatory response have been documented (16–20), we asked whether elevated extracellular MTA could directly impact bacterial virulence. We examined the ability for *S.* Typhimurium pretreated with MTA (300 µM) to induce pyroptosis and invade human cells. As our original studies that identified MTA as a modulator of pyroptosis were performed in lymphoblastoid cells (LCLs), and because B cells are a natural target of *S.* Typhimurium invasion *in vivo* (21–23), we first looked for an effect of exogenous MTA in LCLs. We assessed both pyroptosis and invasion by pairing a modified gentamicin protection assay with flow cytometry, as previously described (24) (Figure 1A). Briefly, pyroptosis was measured by quantifying the number of cells that stained positive for 7AAD three hours post infection with *S.* Typhimurium. Independently, cellular invasion was quantified by using bacteria with a GFP plasmid, inducing GFP production after gentamicin treatment, and quantifying the number of GFP-positive host cells three hours post infection. MTA pretreatment had no effect on bacteria growth from the initial overnight culture dilution through late-log phase to induce SPI-1 gene expression (Figure 1B). MTA pretreated bacteria displayed a 30% decrease (p=0.0001) in their ability to induce pyroptosis (Figure 1C). Furthermore we observed a 35% decrease (p=0.007) in their ability to invade LCLs (Figure 1C). We observed similar effects of MTA on pyroptosis and invasion in THP-1 monocytes (Figure 1D). This was in contrast to our previous finding that MTA treatment of host cells primes them to undergo higher levels of pyroptosis upon *S.* Typhimurium infection (16), and suggests that the molecule has separate effects on both the host and pathogen.

**Figure 1:**
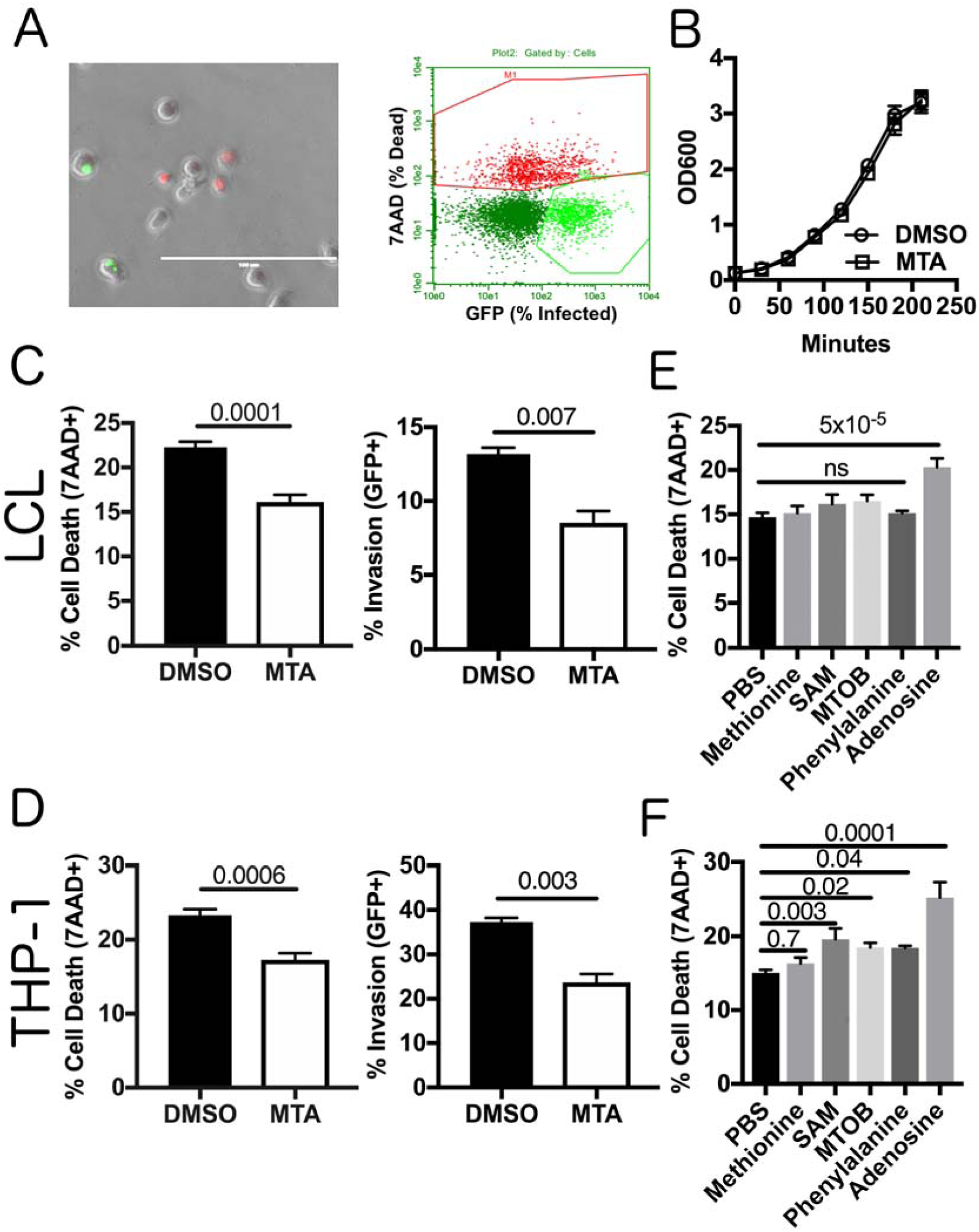
MTA treatment of *S*. Typhimurium reduces pyroptosis and invasion *in vitro*. (A) Modified gentamicin protection assay using an inducible GFP plasmid was used to detect pyroptosis (7AAD+ red nuclear staining) and host cell invasion (intracellular GFP+ bacteria). This is observable by light microscopy and quantifiable by flow cytometry. Scale bar represents 100µm. (B) Exogenous MTA has no effect on growth of *S*. Typhimurium in rich media. OD600 for *S*. Typhimurium treated with 300 µM MTA or 0.5% DMSO were measured every 30 minutes and exhibited equivalent growth (n=3). (C-D) Treatment of bacteria with 300 μM MTA during growth to late log phase (two hours and forty minutes) reduced pyroptosis (MOI 30) and host cell invasion (MOI 10) in LCLs measured three hours post infection in LCL 18592 (C) and THP-1 cells (D). Percent cell death represents all 7AAD+ cells in each infected condition, with baseline uninfected cell death subtracted. See gating in Fig 1A. Data were normalized to the global mean across five experiments and p-values were generated by a student’s t-test. (E, F) Treating bacteria with other methionine related metabolites (methionine, SAM, MTOB, and phenylalanine) or adenosine does not suppress host cell pyroptosis (MOI 30), based on 3-5 independent experiments in LCL 18592 (E) or THP-1 cells (F). Data were normalized to the global mean and p-values were generated by a one-way ANOVA with Dunnett’s multiple comparisons test. All error bars represent the standard error of the mean.

This reduction in pyroptosis could not be reproduced by treating the bacteria with other metabolites related to the mammalian methionine salvage pathway, including methionine, SAM (also known as AdoMet), α-Keto-γ-(methylthio)butyric acid (MTOB), and phenylalanine (Figure 1E, F). Similarly, adenosine (which is only lacking the methylthio group of MTA) was not sufficient to suppress pyroptosis induction. In fact, adenosine increased pyroptosis in both LCLs and THP-1s. Of note, the bacterial cell is reportedly impervious to SAM (25, 26), so we cannot rule out that high intracellular concentrations of the molecule could suppress pyroptosis, however, our results rule out the molecule as an external signal regulating pyroptosis. Thus, these data demonstrate that MTA exposure uniquely suppresses the ability of *S.* Typhimurium to induce pyroptosis and invade host cells.

### *metJ* deletion in *S.* Typhimurium elevates levels of MTA

In order to provide independent evidence for MTA-mediated regulation of *Salmonella* virulence, we genetically disrupted the *S.* Typhimurium methionine metabolism pathway. The protein MetJ is the master repressor of the methionine regulon and transcriptionally blocks multiple enzymatic steps that enable the generation of methionine and SAM (Figure 2A) (10). We hypothesized that deletion of *metJ* would relieve this transcriptional suppression and result in elevated intracellular MTA levels. Therefore, we generated a Δ*metJ* mutant and performed mass spectrometry to examine how metabolites in the methionine metabolism pathway were impacted by the mutation. In line with previous reports, we observed an increase in methionine levels in the Δ*metJ* mutant (27) (Figure 2B). MTA, SAM, and phenylalanine were also increased (Figure 2C, D, E). Expressing metJ from a plasmid could reverse these increases. Consistent with MTA being an inhibitor of polyamine synthesis, the polyamine spermine was decreased in the Δ*metJ* mutant, while no change was observed in another polyamine, spermidine (Figure 2F, G). Disrupting the methionine metabolism pathway by deleting *metJ* did not affect bacterial growth (Figure 2H).

**Figure 2:**
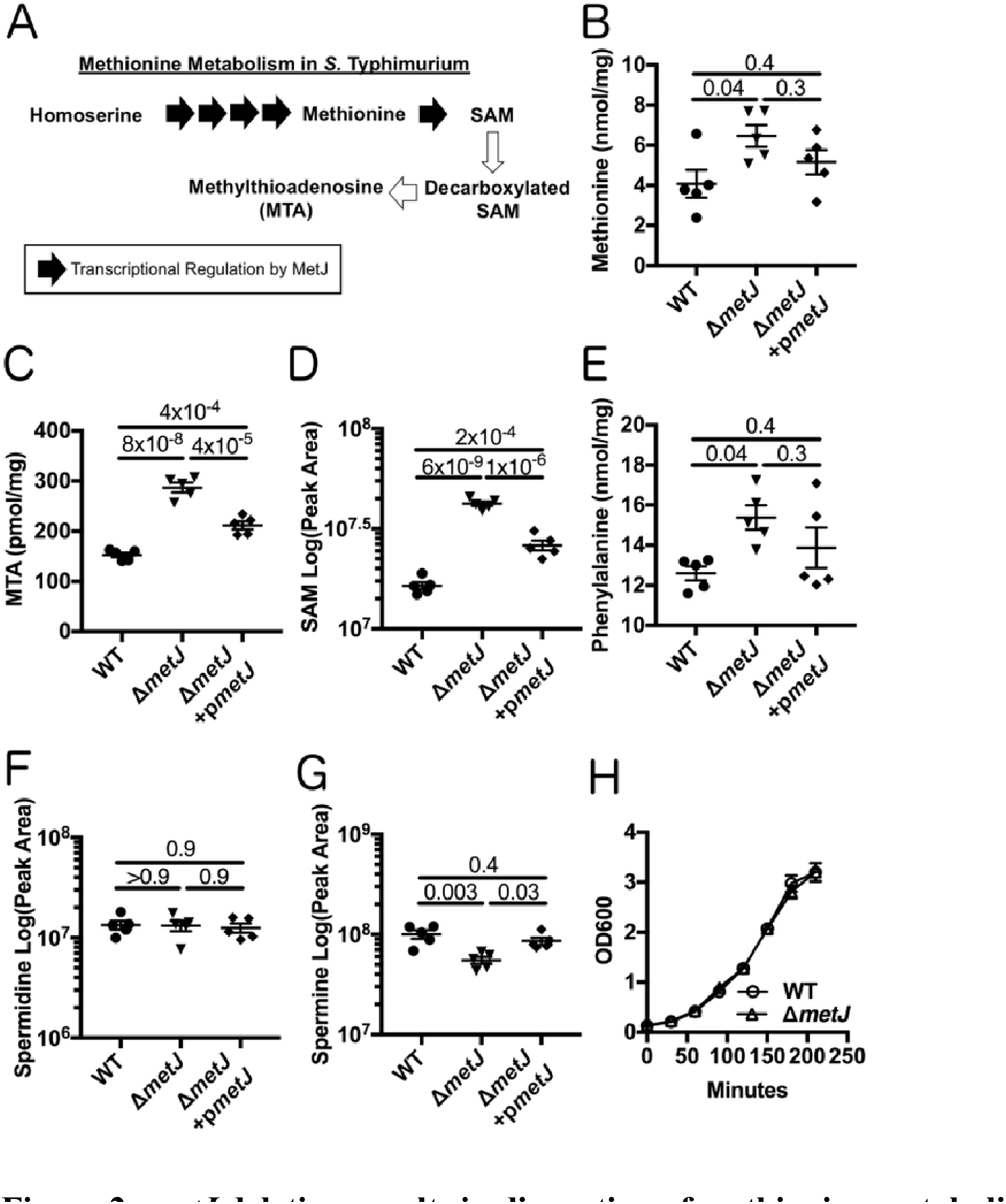
*metJ* deletion results in disruption of methionine metabolism. (A) MetJ regulates the generation of methionine and SAM in *S.* Typhimurium by transcriptionally repressing the methionine regulon. Black arrows represent enzymes transcriptionally repressed by MetJ. Deletion of *metJ* leads to increased methionine (B), MTA (C), SAM (D), phenylalanine (E), as measured by mass spectrometry (n=5 biological replicates). *metJ* deletion did not affect spermidine concentrations (F), but did result in decreased spermine (G). P-values were generated through a one-way ANOVA with Tukey’s multiple comparison test. (H) *metJ* deletion did not affect bacterial growth in LB (n=3 biological replicates, grown in LB+0.5% DMSO). All error bars represent the standard error of the mean.

### Elevated endogenous MTA suppresses pyroptosis and invasion in vitro

After demonstrating that *metJ* deletion leads to the accumulation of MTA in the bacterial cell, we examined whether critical *Salmonella* virulence processes are suppressed in the Δ*metJ* mutant. Similar to *S.* Typhimurium treated with MTA, Δ*metJ* had a reduced ability to induce pyroptosis in LCLs and THP-1 monocytes (Figure 3A, B). Importantly, MTA was the only measured elevated metabolite in the Δ*metJ* mutant that inhibited levels of pyroptosis when added exogenously to wild-type bacteria (see Figure 1D). This reduction in pyroptosis was rescued by expressing *metJ* from a plasmid. Δ*metJ S*. Typhimurium had reduced invasion of LCLs, THP-1 monocytes, and HeLa cells (Figure 3A, B, C). Spermine and other polyamines could not rescue pyroptosis induced by Δ*metJ S.* Typhimurium, suggesting these findings are not mediated by the reduction in spermine we observe in the Δ*metJ* mutant (Figure 3D, E).

**Figure 3:**
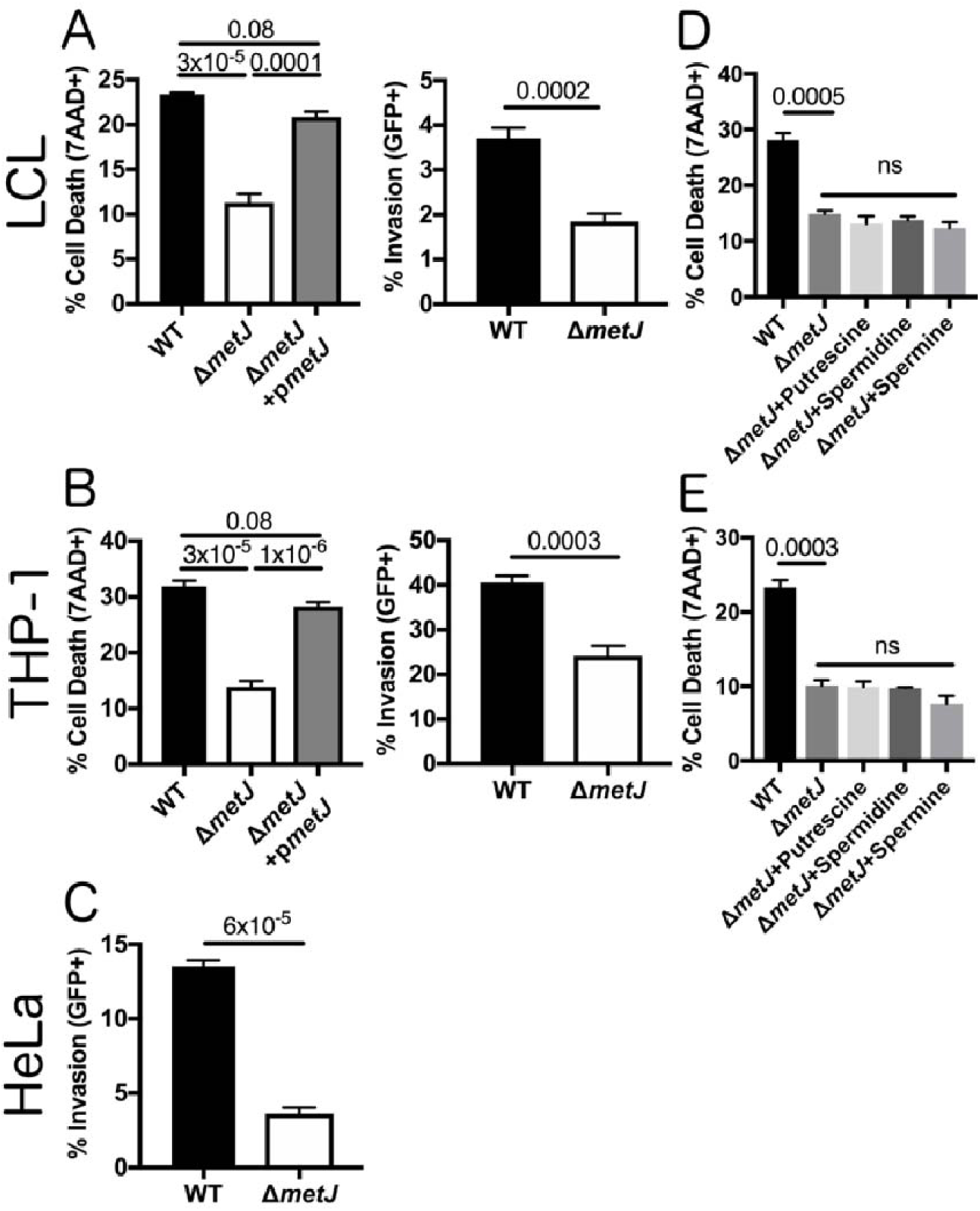
*metJ* deletion reduces pyroptosis and invasion in vitro. (A) Deletion of *metJ* reduces pyroptosis and invasion of 18592 LCLs. (B) Deletion of *metJ* reduces pyroptosis and invasion of THP-1 monocytes. (C) Deletion of metJ reduces invasion of HeLa cells. For A, B, and C, pyroptosis and invasion were measured three hours post infection using a modified gentamicin protection assay across at least three independent experiments. Percent cell death represents all 7AAD+ cells in each infected condition, with baselin uninfected cell death subtracted. See gating in Fig 1A. Data were normalized to the global mean, and p-values were calculated either through a one-way ANOVA with Tukey’s multiple comparisons test or by a student’s t-test. (D, E) Suppression of pyroptosis could not be rescued by treating bacteria with polyamines. Bacteria were treated with 300μM of putrescine, spermidine, or spermine two hours and forty minutes prior to infection to determine whether the effects on pyroptosis were due to effects on polyamine synthesis. For all experiments with 18592 LCLs or THP-1s, cells were infected at MOI 30. HeLas were infected at MOI 5. For D and E, data were generated from two independent experiments normalized to the global mean, and p-values were generated by a one-way ANOVA with Dunnett’s multiple comparisons test. Error bars represent the standard error of the mean.

Motility and the SPI-1 type III secretion system (T3SS) are critical processes for the induction of pyroptosis and *S.* Typhimurium invasion *in vitro*. Motility increases the frequency by which interactions between host and *S.* Typhimurium cells occur and enables bacterial scanning of the host cell surface to optimize invasion (28, 29). The SPI-1 secretion system not only enables the transport of effector proteins into the host cell that enable invasion (1, 2), but also acts as a trigger for pyroptosis in human cells (30–34). Therefore, our observation that both pyroptosis and invasion are suppressed in MTA-treated and Δ*metJ S.* Typhimurium led us to hypothesize that motility and/or SPI-1 are suppressed in response to increased concentrations of MTA.

### Disruption of methionine metabolism and treatment with exogenous MTA impairs *S*. Typhimurium motility

In order to determine whether motility was impaired in the Δ*metJ* mutant, we performed a standard bacterial soft agar motility assay (35) and found that Δ*metJ* was only able to traverse two thirds the distance of the wild-type bacteria over six hours (Figure 4A). This motility defect was restored by expression of *metJ* from a plasmid. In order to confirm that increased MTA levels were sufficient to drive this phenotype, we also examined *S.* Typhimurium motility on soft agar containing 300µM MTA. Like Δ*metJ*, wild-type bacteria swimming on MTA containing agar demonstrated approximately two thirds the motility of those on DMSO (Figure 4B). To further characterize this motility defect, we examined expression of key motility regulators by qPCR in the Δ*metJ* mutant. While the master regulator to the flagellar regulon, *flhD*, did not show reduced expression, two class two flagellar genes, *fliA* and *fliZ,* showed significant downregulation, which correlates with the decrease in motility (Figure 4C). Together, these data revealed that MTA suppresses the *S.* Typhimurium flagellar regulon downstream of *flhD* transcription, resulting in impaired motility *in vitro*.

**Figure 4:**
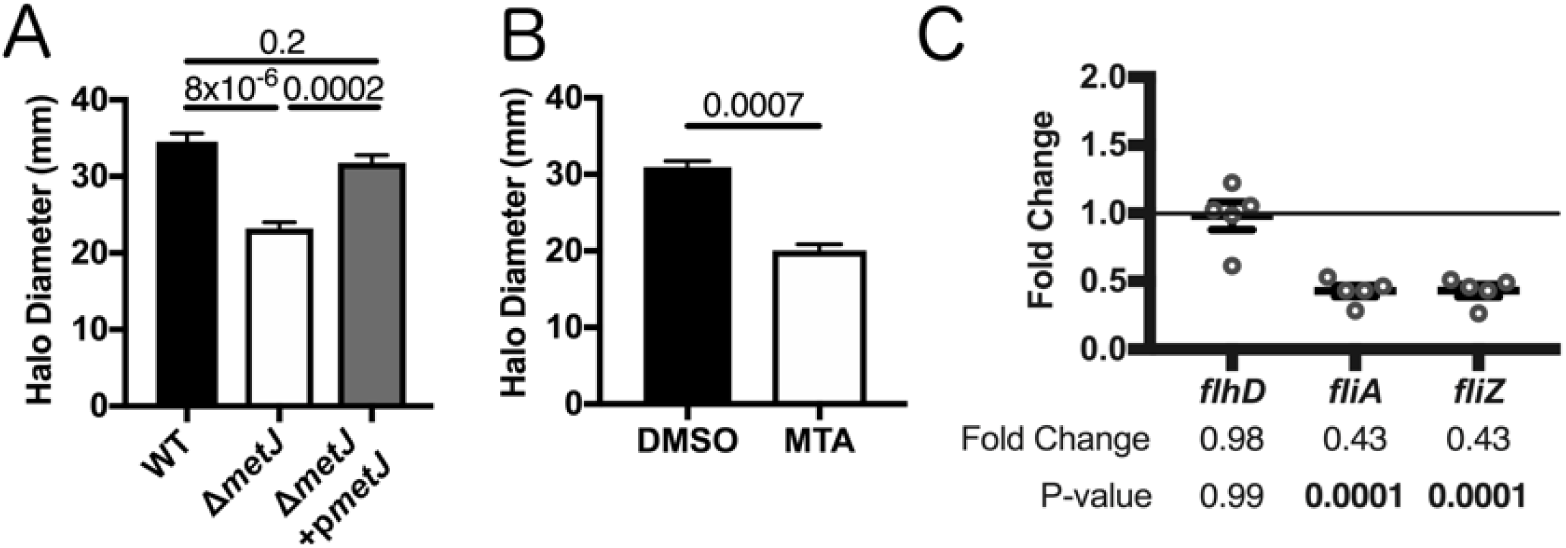
*metJ* deletion and MTA treatment of *S.* Typhimurium reduces motility. (A) *S.* Typhimurium motility is suppressed in Δ*metJ* mutants. Motility was measured after six hours at 37°C on 0.3% LB agar. (B) Exogenous MTA suppresses *S.* Typhimurium motility. Motility was measured after six hours on 0.3% LB agar with 0.5% DMSO or 300μM MTA. Data from B and C represent at least three independent experiments normalized to the global mean, and p-values were calculated by a one-way ANOVA with Tukey’s multiple comparisons test or a student’s t-test. (C) Flagellar genes are suppressed in Δ*metJ*. RNA was extracted from wild-type *S.* Typhimurium or Δ*metJ* mutant bacterial cultures grown to late log phase in LB broth (n=5) and analyzed by qPCR with the ribosomal *rrs* gene serving as the endogenous control. Each dot represents an independent biological replicate. P-values were calculated by a one-way ANOVA with Dunnett’s multiple comparisons test.

### Elevated endogenous MTA suppresses expression of SPI-1 encoded genes

Deploying the SPI-1 encoded T3SS depends on a complex regulatory network in which HilD, HilC, and RtsA drive expression of *hilA* (36, 37). HilA then enables the expression of the *inv/spa* and *prg/org* operons (38–40). Both HilD and HilA directly promote the expression of the first gene in the *inv/spa* operon, *invF* (41), which is then able to drive the expression of the *sic/sip* operon and a number of other critical effectors (39, 42, 43).

Three lines of evidence demonstrated suppression of SPI-1 by MTA. First, we observed suppression of SPI-1 encoded genes in Δ*metJ S.* Typhimurium by qPCR. In particular, we saw a decrease in *invF* expression, as well as the translocon component *sipB*. The finding that *sipB* is significantly suppressed is of note, as SipB is involved in induction of pyroptosis by *S.* Typhimurium (33). Further, while not statistically significant when taking into account multiple-test correction, we saw comparable changes trending towards significance in the regulatory factor *rtsA*, the chaperone protein *sicP*, and the needle complex component *prgH*. (Figure 5A). This suggests that there is at least modest suppression of SPI-1 regulated genes by MTA on the transcriptional level under standard culture conditions. Second, reduced expression of SipA, a SPI-1 secreted effector, was detected in Δ*metJ* cell lysates by western blot staining (Figure 5B). Third, this reduction in SipA was greater in Δ*metJ* cell free supernatants than cell lysates, suggesting that both expression of SipA as well as its secretion by the T3SS apparatus are suppressed in the mutant (Figure 5B). This is consistent with the suppression *of invF* and *sipB* observed by qPCR. These reductions in SipA were also detected in MTA treated *S.* Typhimurium (Figure 5C). Together, these data demonstrate suppression of the SPI-1 network by MTA.

**Figure 5:**
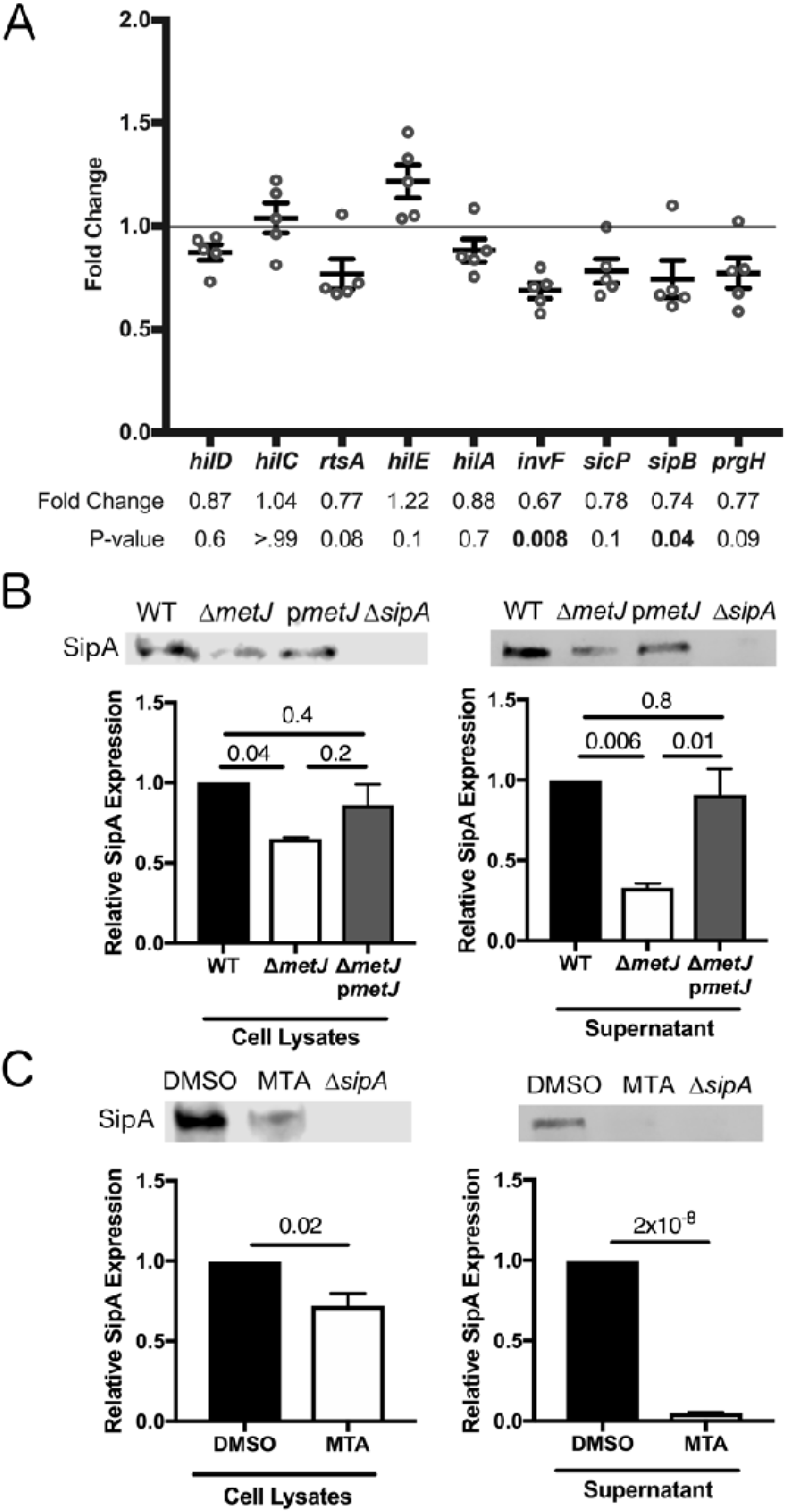
*metJ* deletion and MTA treatment of *S.* Typhimurium reduces SPI-1 secretion. (A) SPI-1 genes are suppressed in Δ*metJ* mutants. RNA was extracted from wild-type *S*. Typhimurium or Δ*metJ* mutant bacterial cultures grown to late log phase in LB broth (n=5) and analyzed by qPCR with the ribosomal *rrs* gene serving as the endogenous control. Each dot represents an independent biological replicate. P-values were calculated by a one-way ANOVA with Dunnett’s multiple comparisons test. (B) SipA secretion is suppressed in Δ*metJ* mutants. SipA protein was measured by western blotting cell lysates or cell free supernatants collected at late log phase growth. (C) SipA secretion is suppressed in *S.* Typhimurium treated with exogenous MTA. SipA protein was measured by western blotting cell lysates or cell free supernatants collected at late log phase growth from bacteria grown either in 0.5% DMSO or 300μM MTA. For E and F, cell lysates were normalized by total protein content. Cell free supernatants were spiked with 100ng/μL of BSA as a loading control, concentrated by TCA acid precipitation, and normalized by volume and total protein. Data represent three independent experiments and are normalized to wild-type expression. P-values were calculated by a one-way ANOVA with Tukey’s multiple comparisons test or a student’s t-test. All error bars represent the standard error of the mean.

### Disruption of bacterial methionine metabolism impairs virulence *in vivo*

Induction of inflammation and host cell invasion are critical processes that enable *Salmonellae* to colonize and disseminate from the mouse gut (2, 8, 36, 44–46). Our findings that both the induction of pyroptosis and host cell invasion are attenuated in the Δ*metJ* mutant led us to hypothesize that this mutant also has impaired virulence *in vivo.* To test this, we orally infected C57BL/6J mice with wild-type and Δ*metJ S.* Typhimurium and measured the bacteria’s ability to infect and disseminate to the spleen. We found that Δ*metJ* had a 30-fold reduction in fitness compared to wild-type *S.* Typhimurium at five days post infection (Figure 6). This supports the hypothesis that metJ is important for *S.* Typhimurium to establish an infection in the mammalian gut and disseminate to the spleen.

**Figure 6:**
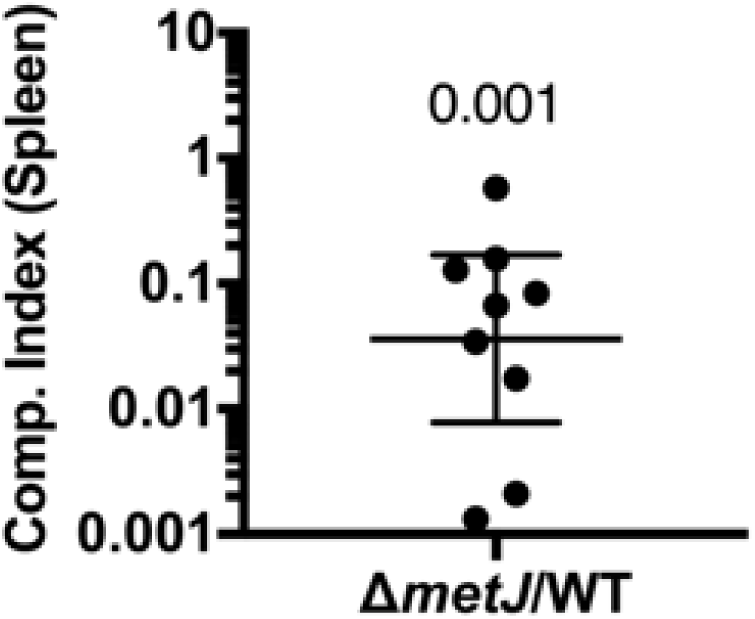
*metJ* deletion suppresses *S.* Typhimurium virulence *in vivo*. *metJ* deletion reduced bacterial fitness in oral models of infection. C57BL/6J mice were infected with 10^6^ total *S.* Typhimurium from a 1:1 mixture of wildtype and Δ*metJ* bacteria by oral gavage. Spleens were harvested five days post infection and bacteria quantified to calculate the competitive index. Competitive index from each mouse is graphed as (Δ*metJ* CFUs/WT CFUs)/(Δ*metJ* CFUs in inoculum/WT CFUs in inoculum). P-value was calculated by log transforming these ratios and comparing to an expected value of 0 using a one-sample t-test. Data are from three independent experiments, and are graphed using the geometric mean and 95% confidence interval.

### Disruption of methionine metabolism in *S.* Typhimurium reduces inflammatory cytokine production

We previously reported that treatment of mice with MTA before infecting with a lethal dose of *S*. Typhimurium resulted in reduced production of sepsis-related cytokines (IL-6 and TNFα) and modestly prolonged survival (17). We hypothesized that the previously observed effects on inflammation may be due to MTA’s impact on the microbe. This hypothesis is based on our findings that *metJ* knockout suppresses murine-detected PAMPs, specifically flagellin (47, 48) and the SPI-1 T3SS (34, 49, 50).

To test whether MTA reduces host inflammation by suppressing *S.* Typhimurium virulence, we injected mice with a lethal dose (1×10^6^ CFUs) of either wild-type or Δ*metJ S.* Typhimurium by an intraperitoneal route and measured CFUs and cytokines four hours post infection. Similar to what we observed with exogenous treatment of the mice with MTA (17), we did not see a difference in CFUs, suggesting that both wild-type and Δ*metJ S.* Typhimurium were equally capable of colonizing the spleen at this time point (Figure 7A). In contrast, IL-6 was 28% lower (p=0.03) in mice infected with the Δ*metJ* mutant (Figure 7B). Further, TNFα concentrations also showed a similar relative decrease (34%), though that result did not meet statistical significance (p=0.09). Therefore, Δ*metJ* phenocopies the anti-inflammatory effects of treating mice with MTA before infection with *S.* Typhimurium, demonstrating that these effects of MTA could be mediated not only through effects on the host, but also through suppression of *S.* Typhimurium pro-inflammatory virulence gene expression.

**Figure 7:**
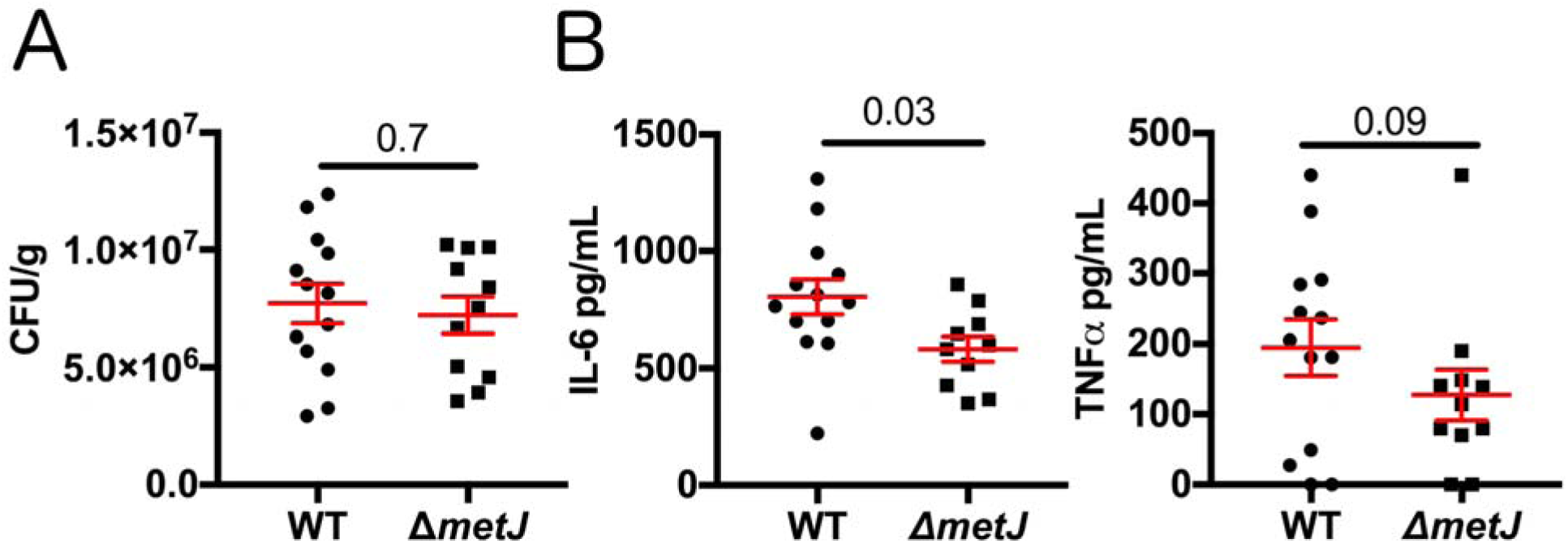
Bacterial methionine metabolism reduces host inflammation. (A) *metJ* deletion reduced bacterial fitness at four hours post intraperitoneal infection. C57BL/6 mice were infected with 10^6^ wild-type or Δ*metJ S.* Typhimurium. Four hours post infection spleens were harvested and CFUs quantified. Data are graphed using the geometric mean and the 95% confidence interval. (B) *metJ* deletion reduced the host cytokine response to *S.* Typhimurium infection. Plasma harvested four hours post infection showed reduced sepsis-associated cytokines, IL-6 and TNFα. Data were generated by ELISA. Error bars represent the standard error of the mean. All data represent three independent experiments normalized to th global mean, with each dot representing a biological replicate. P-values for CFUs and IL-6 were calculated by an unpaired t-test with Welch’s correction. Because the TNFα data are non-normal, a Kolmogorov-Smirnov test was used to calculate the p-value.

## Discussion

Here we report that exposing *S.* Typhimurium to exogenous MTA or increasing endogenous MTA production suppresses virulence *in vitro* and *in vivo*. These phenotypes correlate with transcriptional, translational, and functional suppression of the flagellar regulon and SPI-1. This adds to a growing body of literature demonstrating that environmental factors in the host can regulate critical *Salmonella* virulence factors (3-7, 51, 52). We hypothesize that this represents an example of host-pathogen cross-talk, in which the host suppresses *Salmonella* virulence by increasing MTA concentrations. This would represent a novel antimicrobial mechanism, and is supported by our findings that MTA plasma concentrations increase during infection (17). Future studies could benefit from examining MTA concentrations in the gut, as modulation of MTA there could directly promote or impair SPI-1 function. If MTA were increased in the gut during infection, as we observed in plasma, this could result in suppression of SPI-1 and bacterial invasion. Alternatively, if MTA were present at lower levels in the ileum relative to the rest of the gut, this could serve as a signal to turn on expression of the SPI-1 T3SS. Our future studies will investigate if and how MTA functions as a cross-kingdom signal during *Salmonella* infection and how methionine metabolism can be therapeutically targeted.

While no previous work examined the impact of *metJ* deletion on *S.* Typhimurium virulence, two papers reported effects of Δ*metJ* and virulence in other bacterial pathogens. Bogard *et al*. report that Δ*metJ* suppresses *Vibrio cholerae* virulence *in vivo* (53). Conversely, Cubitt *et al*. demonstrated that Δ*metJ* increased the production of quorum sensing molecules and expression of virulence genes in the potato pathogen *Pectobacterium atrosepticum* (54). However, in both these cases, the metabolic changes responsible for these phenotypes are unknown. We hypothesize that our discovery of the role of *metJ* in regulating intracellular MTA concentrations could help explain these findings. If exogenous MTA can drive these phenotypes, similar to what we report here in *S.* Typhimurium, it would suggest that modulation of MTA concentrations represents a mechanism by which virulence can be manipulated across multiple bacterial species.

One mechanism by which MTA may influence the *S.* Typhimurium flagellar regulon and SPI-1 encoded genes is by altering methylation. In prokaryotic and eukaryotic systems, SAM provides the methyl for a variety of DNA, RNA, and protein methylation reactions (55–60). In eukaryotic systems, modulation of methionine metabolism resulting in changes to the cellular MTA and SAM pools can have important consequences on protein, DNA, and RNA methylation (61–64). Therefore, we hypothesize that increased MTA leads to altered methylation of critical SPI-1 and flagellar regulators, resulting in their suppression. This could be part of a metabolic stress response, as both SPI-1 secretion and swimming motility are energetically expensive processes that the bacteria may downregulate in response to methionine metabolism dysregulation (65). Understanding how MTA is sensed, what process it regulates directly, and how this process influences virulence could help understand this novel example of host-pathogen communication and further inform future therapeutics targeting this process.

Since exogenous and endogenous MTA affects *Salmonella* virulence, both bacterial and host methionine metabolism present therapeutic targets. Previous studies tested MTA nucleosidase inhibitors against bacterial pathogens based on the assumption that disrupting MTA nucleosidase would lead to an MTA accumulation, resulting in an arrest of cellular growth and reduced bacterial viability (66–68). However, MTA nucleoside inhibitors showed, at most, modest bacteriostatic potential in these studies. In contrast, studies examining the effects of these compounds on quorum sensing also showed no changes in bacterial growth, but did identify suppression of AI-2 synthesis (69, 70). This is in line with our observation that *Salmonella* growth is not impaired by increased concentrations of MTA in the Δ*metJ* mutant, but that there are functional consequences on virulence. However, no study has examined the potential of these compounds to directly impact virulence, independent of growth. Our data suggest that these compounds likely have antibacterial properties, because their disruption of methionine metabolism *in vivo* impairs virulence. Furthermore, other groups have developed S-methyl-5’-thioadenosine phosphorylase (MTAP) inhibitors, which block mammalian MTA catabolism (71–73), increasing MTA concentrations in tissues, plasma, and urine in murine models (74). Based on these results and our demonstration that high extracellular MTA suppresses virulence, we hypothesize that MTAP inhibitors could be a host-directed therapy during *Salmonella* infection. Therefore, the findings described here suggest that MTA nucleosidase inhibitors and MTAP inhibitors could be harnessed to combat bacterial infections and improve clinical outcomes.

## Materials and Methods

### Mammalian cells and bacterial strains

HapMap LCLs were purchased from the Coriell Institute. LCLs and THP-1 monocytes were cultured at 37°C in 5% CO_2_ in RPMI 1650 media (Invitrogen) supplemented with 10% FBS, 2 μM glutamine, 100 U/mL penicillin-G, and 100 mg/mL streptomycin. HeLa cells were grown in DMEM media supplemented with 10% FBS, 1mM glutamine, 100 U/mL penicillin-G, and 100mg/mL streptomycin. Cells used for *Salmonella* gentamicin protection assays were grown in antibiotic free media one hour prior to infection.

All *Salmonella* strains are derived from the *S.* Typhimurium NCTC 12023 (ATCC 14028) strain and are listed in Table 1. All knockout strains were generated by lambda red recombination (75). Recombination events were verified by PCR, and the pCP20 plasmid was used to remove the antibiotic resistant cassette after recombination (76). Strains were cultured overnight in LB broth (Miller) overnight, subcultured 1:33, and grown for two hours and forty minutes shaking at 37°C before all experiments unless otherwise noted. Strains containing the temperature sensitive plasmids pKD46 or pCP20 were cultured at 30°C and removed at 42°C. Ampicillin was added to LB at 50μg/mL, kanamycin at 20μg/mL, and tetracycline at 12μg/mL.

**Table 1:** Bacterial strains used in this study

| **Bacterial Strains** |  |  |  |
| --- | --- | --- | --- |
| **Strain** | **Genotype** | **Plasmid** | **Resistance** |
| DCK543 | NCTC 12023 (ATCC 14028) |  |  |
| DCK545 | $\Delta metJ$ | | |
| DCK546 | NCTC 12023 (ATCC 14028) | pWSK29 | Amp |
| DCK547 | $\Delta metJ$ | pWSK29 | Amp |
| DCK548 | $\Delta metJ$ | pWSK29::*metJ* | Amp |
| DCK571 | NCTC 12023 (ATCC 14028) | *p67GFP* | Amp |
| DCK573 | $\Delta metJ$ | *p67GFP* | Amp |
| DCK574 | NCTC 12023 (ATCC 14028) | pWSK129 | Kan |
| DCK576 | $\Delta metJ$ | pWSK129 | Kan |
| DCK88 | sspA::Tn5 (*sipA* insertion) |  | Tet |

Exogenous metabolites (5′-Deoxy-5′-(methylthio)adenosine, L-Phentlalanine, L-Methionine, α-Keto-γ-(methylthio)butyric acid sodium salt, Adenosine, Spermine, Spermidine, Putrescine dihydrochloride: Sigma. S-(5’-Adenosyl)-L-methionine chloride (hydrochloride): Cayman Chemicals) were added to LB during the two hour and forty minute subculture step at 300μM final concentration unless otherwise noted.

### Gentamicin protection assay

As previously described, inducible GFP plasmids were transformed into *S.* Typhimurium strains in order to assess both *Salmonella*-induced cell death and invasion by flow cytometry (24). Briefly, bacteria cultures were prepared as described above and used to infect LCLs and THP-1 monocytes (MOI 10 or MOI 30), as well as HeLa cells (MOI 5). At one hour post infection cells were treated with gentamicin (50μg/mL), and IPTG was added 2 hours post infection to induce GFP expression. At 3.5 hours post infection cells were assessed for cell death using either staining with 7-aminoactinomycin D (7-AAD; Biomol) read by a Guava Easycyte Plus flow cytometer (Millipore). Percent invasion was determined by quantifying the number of GFP+ cells 3.5 hours post infection using the Guava Easycyte Plus flow cytometer.

### Metabolomics

Bacteria were grown overnight as described above and subcultured 1:33 in 10mL LB and grown for 2 hours and 40 minutes. After thorough washing in PBS, samples were flash frozen, thawed, and 0.5 mL PBS was added directly onto the pellets. Samples were then transferred to 2 mL CK01 bacterial lysis tubes (Bertin). These were then taken through 3 cycles of 20 second bursts at 7,500 RPM with 30 second pauses in between bursts using a Bertin Precellys (protocol as recommended by Bertin). Samples were spun at 5,000 for 5 minutes and a Bradford assay was performed on each lysate to gather protein concentration values. 100 µL from each homogenate was pipetted directly into a 2 mL 96-well plate (Nunco).

The internal standard methanol solution was made by pipetting 166.7 µL of NSK-A standard (Cambridge Isotope) at 500 µM, 62.5 µL of 500 nM of d3-MTA, and 49.771 mL of MeOH. 900 µL of this internal standard solution in MeOH was pipetted into all of the standard and sample wells. The plate was then capped and mixed at 700 rpm at 25C for 30 minutes. The plate was then centrifuged at 3000 rpm for 10 minutes. Using an Integra Viaflo96 pipettor, 600 µL of extract was pipetted out and transferred to a new 96-well plate. The extracts were allowed to dry under a gentle stream of nitrogen until completely dry. 32 µL of 49/50/1 water/acetonitrile/trifluoroacetic acid was added to each well and mixed at 650 rpm for 10 minutes at room temperature. Then 128 µL of 1% trifluoroacetic acid was added to each, mixed briefly, and centrifuged down to give a total of 160 µL of sample.

The samples were analyzed using Ultraperformance Liquid Chromatography/Electrospray Ionization/Tandem Mass Spectrometry (UPLC/ESI/MS/MS) using a customized method allowing chromatographic resolution of all analytes in the panel. Flow from the LC separation was introduced via positive mode electrospray ionization (ESI+) into a Xevo TQ-S mass spectrometer (Waters) operating in Multiple Reaction Monitoring (MRM) mode. MRM transitions (compound-specific precursor to product ion transitions) for each analyte and internal standard were collected over the appropriate retention time. The data were imported into Skyline (https://skyline.gs.washington.edu/) for peak integration, and exported into Excel for further calculations.

### Bacterial RNA isolation and qPCR

Bacteria were grown as described above and RNA was isolated from 5×10^8^ bacteria using the Qiagen RNAprotect Bacteria Reagent and RNeasy minikit (Qiagen) according to the manufacturer’s instructions. RNA was treated with DNase I (NEB) and 500ng were reverse transcribed using the iScript cDNA synthesis kit (Bio-Rad Laboratories). qPCR was performed using the iTaq Universal SYBR Green Supermix (Bio-Rad Laboratories). 10μL reactions contained 5μL of the supermix, a final concentration of 500nM of each primer, and 2μL of cDNA. Reactions were run on a StepOnePlus Real-Time PCR System (Applied Biosystems). The cycling conditions were as follows: 95°C for 30 seconds, 40 cycles of 95 degrees for 15 seconds and 60°C for 60 seconds, and 60°C for 60 seconds. A melt curve was performed in order to verify single PCR products. The comparative threshold cycle (C_T_) method was used to quantify transcripts, with the ribosomal *rrs* gene serving as the endogenous control. ΔC_T_ values were calculated by subtracting the C_T_ value of the control gene from the target gene, and the ΔΔC_T_ was calculated by subtracting the wildtype ΔC_T_ from the mutant ΔC_T_ value. Fold change represents 2^-ΔΔCT^. Experiments included three technical replicates and data represent qPCR results from five separate RNA isolation experiments. Oligonucleotides are listed in Table 2.

**Table 2:**
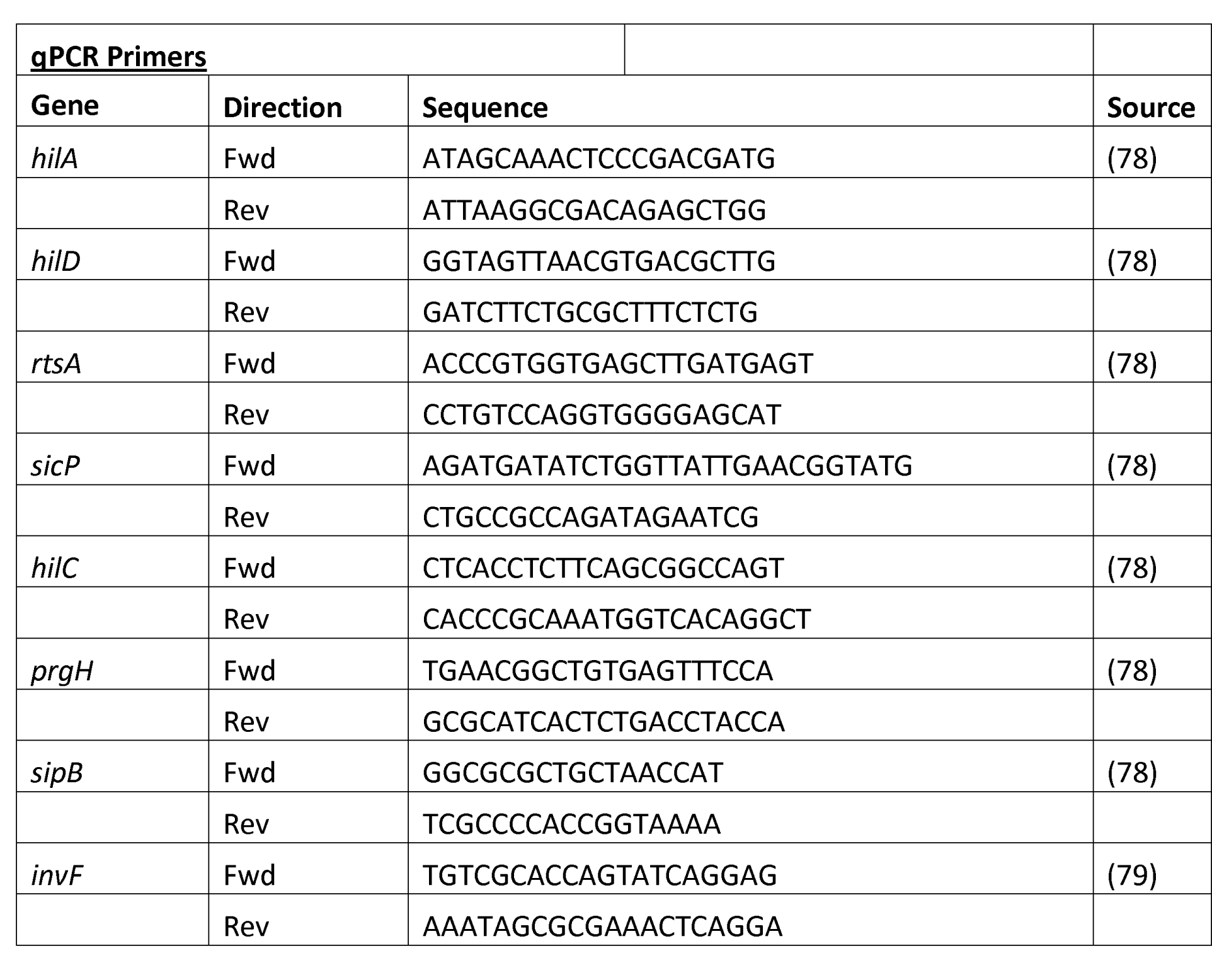
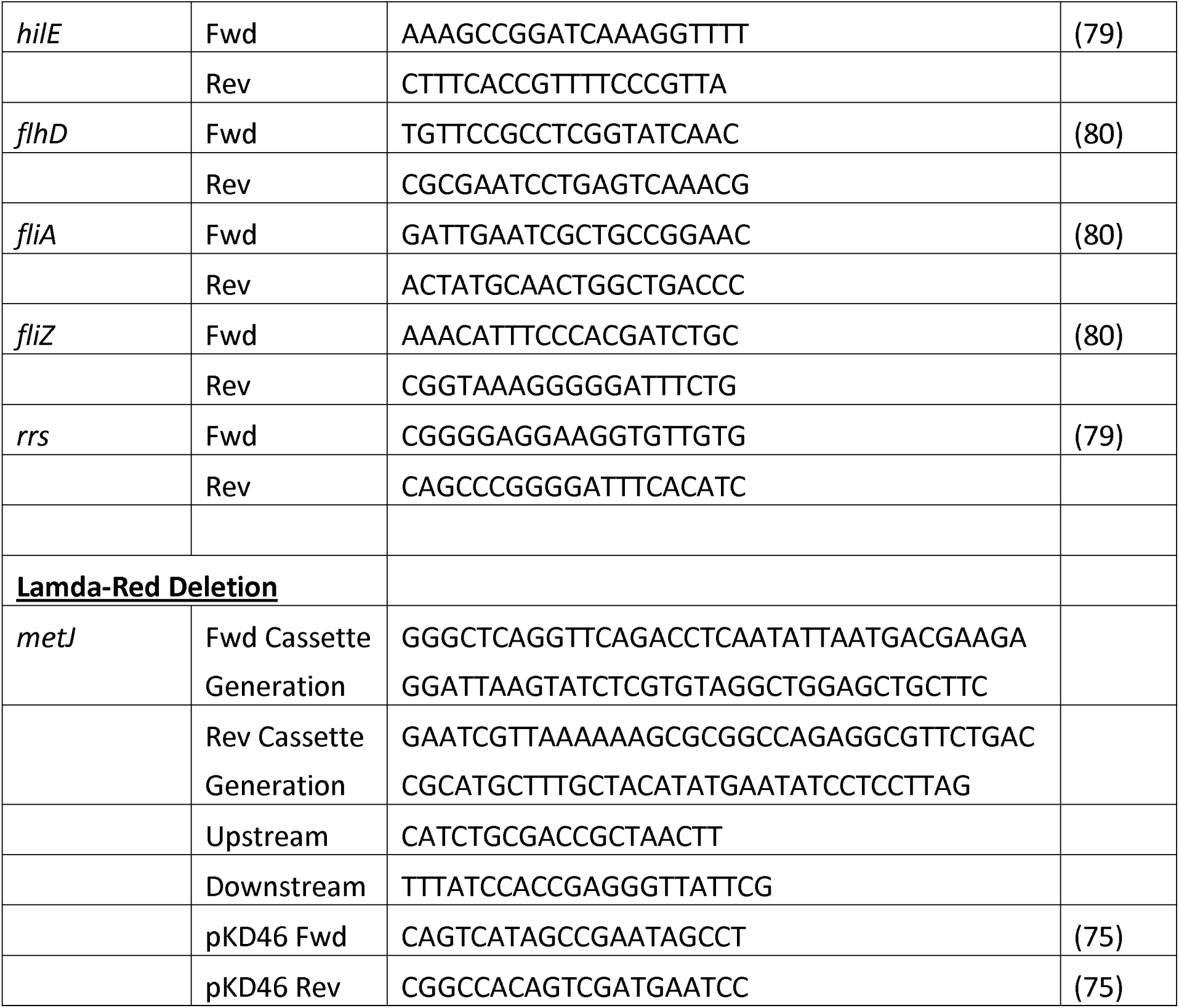
Oligonucleotides used in this study

### Analysis of Bacterial Protein Expression

Bacteria were grown as described above. For analysis of cell lysates, bacterial cultures were centrifuged at 10,000xg for 5 minutes. Supernatant was discarded and pellets were lysed in 2x laemmli buffer (Bio-Rad) with 5% 2-Mercaptoethanol. Samples were boiled for 10 minutes and analyzed on Mini-PROTEAN TGX Stain-Free gels (Bio-Rad). Bands were stained with a rabbit anti-SipA antibody overnight at 4°C. Antibody were then detected by staining with the LI-CORE IRDye 800CW donkey anti-rabbit antibody. SipA was quantified using a LI-CORE Odyssey Fc, paired with Image Studio software. Bands were quantified to total protein using the TGX stain free system. Total protein was quantified with using Fiji (77).

For secreted protein analysis, cultures were centrifuged at 10,000xg for 5 minutes, and supernatants were passed through a .2μm syringe filter. At this point, 6μL of 100ng/μL BSA was added to 600μL of supernatant as a loading control. Chilled 100% trichloroacetic acid was added to a final concentration of 10% and incubated on ice for 10 minutes. 600μL of chilled 10% trichloroacetic acid was added, and the solution incubated on ice for another 20 minutes before being centrifuged at 20,000xg for 30 minutes. Pellets were washed twice with acetone and resuspended in 2x laemmli buffer (Bio-Rad) with 5% 2-Mercaptoethanol before boiling for 10 minutes. Proteins were then analyzed as described above.

### Motility Assay

Motility assay was performed as previously described (35). Briefly, strains were cultured overnight in LB broth (Miller), subcultured 1:33, and grown for two hours and forty minutes shaking at 37°C. 2μL of the subcultured solution was plated in the center of a 0.3% agar LB plate supplemented with 50μg/mL ampicillin. Metabolites or DMSO were added to the solution prior to the agar solidifying in order to allow exposure of the bacteria to the metabolite for the entirety of the assay. Plates were incubated at 37°C for 6 hours before the halo diameter was quantified.

### Mouse infection studies

Mouse studies were approved by the Duke Institutional Animal Care and Use Committee and adhere to the *Guide for the Care and Use of Laboratory Animals* of the National Institutes of Health. Bacteria were grown as described above, washed, and resuspended in PBS. Inoculums were confirmed by plating for CFUs. For oral infections, age and sex matched 7-16 week old C57BL/6J mice were fasted for 12 hours before infection. Thirty minutes prior to infection mice received 100μL of a 10% sodium bicarbonate solution by oral gavage. Age and sex matched mice were then infected with 10^6^ bacteria in 100μL PBS by oral gavage. Five days post infection, mice were euthanized by CO_2_ asphyxiation, and spleens were harvested, homogenized, weighed, and plated on LB agar containing either ampicillin or kanamycin. Competitive index was calculated as (Δ*metJ* CFUs/WT CFUs)/(Δ*metJ* CFUs in inoculum/WT CFUs in inoculum). Statistics were calculated by by log transforming this ratio from each mouse and comparing to an expected value of 0 using a one-sample t-test.

For high dose intraperitoneal (IP) injection, cultures were grown as described above, washed, and resuspended in PBS. Inoculums were confirmed by plating for CFUs. Age and sex matched 6-8 week old C57BL/6J mice then received 10^6^ bacteria in 100μL of PBS by IP injection. Four hours post infection, mice were euthanized by CO_2_ asphyxiation, blood was drawn by cardiac puncture, and spleens were harvested, homogenized, weighed, and plated on LB agar containing ampicillin. Plasma was isolated using PST tubes with lithium heparin (BD). IL-6 and TNFα were then quantified from plasma and spleen extracts using the DuoSet ELISA kits (R&D Systems).

## Acknowledgements

We thank David W. Holden for providing an intellectual home for JJG during the formative part of this project and for providing useful discussion of the manuscript. We also thank Kyle Gibbs, Monica Alvarez, Alejandro Antonia, Sarah Jaslow, and Kelly Pittman for sharing their expertise and support throughout the project. JSB was supported by NIH 5T32GM007754. JSB and DCK were supported by NIH R01AI118903 and Duke MGM startup funds. JJG was supported by a Wellcome Trust Clinical PhD Fellowship (102342/Z/13/Z). TLMT was supported by an Imperial College Junior Research Fellowship (RSRO_P50016). We thank the Duke University School of Medicine for the use of the Proteomics and Metabolomics Shared Resource, which provided measurement of MTA and related metabolites.

